# Translation and cross-cultural adaptation to Portuguese of The Patient- And Nutrition-Derived Outcome Risk Assessment Score (PANDORA)

**DOI:** 10.1101/584078

**Authors:** Juliana B. de Lima, Marina B. Campos, Lays S. Ribeiro, Maria I. S. Taboada

## Abstract

**Introduction:** Hospital malnutrition presents alarming rates and is characterized as an independent risk factor for mortality. Hospital mortality has been studied as an important indicator of the quality of care. In this sense, the Patient- And Nutrition-Derived Outcome Risk Assessment Score (PANDORA) was created, seeking to associate the nutritional status and in-patients’ illness data with the risk of death within 30 days. The study aimed to perform the translation, cross-cultural adaptation to Portuguese and application of an instrument of identification of mortality risk in the hospital setting.

**Methods:** A crosssectional study was carried out in a university hospital in the city of Goiania-GO, Brazil, in 2018. A translation and adaptation of the PANDORA instrument was carried out and it was applied to hospitalized patients to evaluate their power to predict mortality.

**Results:** Fifty-four 54 patients were included in the study, most of them female and 33% elderly. More than 16% of the sample presented low weight, which was positively associated with the occurrence of death. The prevalence of cancer was almost 80% and all patients who died had cancer. In the adjusted logistic regression analysis, it was verified that there was no association between the PANDORA score and death in hospitalized patients, however, there was a trend of association of sex and body mass index with death in these patients.

**Conclusions:** In this study, the PANDORA score was not able to predict death in the patients in our sample, but found significant association of low weight at admission with mortality. Further studies are needed for the validation of PANDORA in Portuguese.

## Introduction

Hospital malnutrition has alarming rates despite therapeutic advances, especially in emerging and industrialized countries. It affects almost 50% of adult patients in Latin American countries, including Brazil. About 40% of in-patients are affected by this condition, which represents an independent risk factor for mortality, besides favoring complications during hospitalization [1,2]. The influence of nutritional status on patient prognosis has been reported in the literature some time ago. Correia and Waitzberg (2003) found a 12.4% mortality rate in malnourished patients, three times higher than those considered well-nourished in the study (4.7%), showing that malnutrition is an independent risk factor for the increase in hospital mortality [3].

According to Tsaousi et al. (2014), inadequate feeding can increase up to eight times the risk of hospital mortality, in addition to prolonged hospital stay [4]. This condition is often neglected and presents a high risk of developing other complications, such as surgical and infectious, pressure lesions, increase in length of hospitalization and depletion of the immune system. Its early identification is important to establish the most appropriate nutritional management aiming at better outcomes in these patients [5].

Hospital mortality has been identified for many years as an important indicator of the quality of care provided and has been extensively studied through the application of predictive instruments. One of the first predictive models of death within 30 days of hospital stay was for elderly patients with acute myocardial infarction [6]. These predictive models have been extensively used in emergencies and specific acute situations such as cerebral vascular accident (CVA), acute coronary syndrome (ACS), heart failure (HF), among others, in order to evaluate the quality of care provided [7].

In this sense, Hiesmayr et al. (2015) developed a simple punctuation system to predict mortality in 30 days of hospitalized patients, with scores based on nutrition and baseline disease. The instrument was named *The Patient- And Nutrition-Derived Outcome Risk Assessment Score* (PANDORA), and seeks to associate nutritional status and in-patient disease data with the risk of death within 30 days [2].

From the above, evaluating the risk of hospital death related to nutritional status through a standardized questionnaire is important to assess the effectiveness of services provided in the hospital environment, and may contribute to the establishment of more effective therapeutic plans. Thus, this study aimed to perform the translation, cross-cultural adaptation to Portuguese and application of a risk identification instrument for mortality in the hospital setting.

## Methods Study design

A cross-sectional study was developed in a tertiary university hospital, through the application of the translated and adapted questionnaire.

### Population

The study sample consisted of adult patients hospitalized with any pathologies. Patients older than 18 years, hospitalized in the Medical, Surgical, Tropical or Emergency Clinics were included in the study. Patients under 18 years of age, in intensive care and pregnant patients were excluded.

### Ethical aspects

The study was submitted to the HC-UFG / EBSERH Research Ethics Committee, approved under No. 2,674,012. All participants were informed about the content of the study and the risks involved, by means of a signed Free and Informed Consent Form.

### Procedures for cross-cultural translation and adaptation adopted in the study

The methodology for translation and cross-cultural adaptation of the questionnaire adopted in the study was based on the procedures suggested by Beaton et al. [8], American Educational Research Association, American Psychological Association and National Council on Measurement in Education [9] and reviewed by Muñiz, Elosua and Hambleton [10], according to Fig. 1.

**Fig 1.**
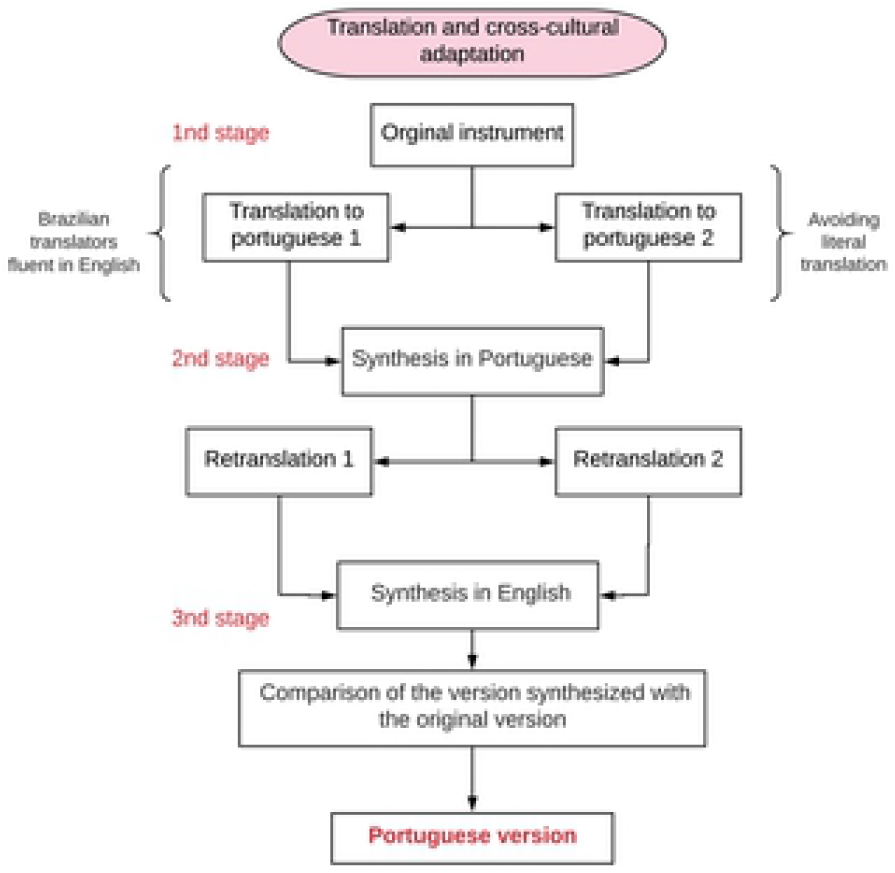
Methodological procedures used in the translation and cross-cultural adaptation of PANDORA into Portuguese

In order to perform the translation and adaptation, the authors of the original instrument gave permission (Fig. 2) for its use, via e-mail. In addition, a committee was formed by the authors of the new version of PANDORA, to discuss concepts adjacent to the adapted test, considering the particularities of the target population. All translators in the study were unaware of the test to be adapted.

**Fig. 2.**
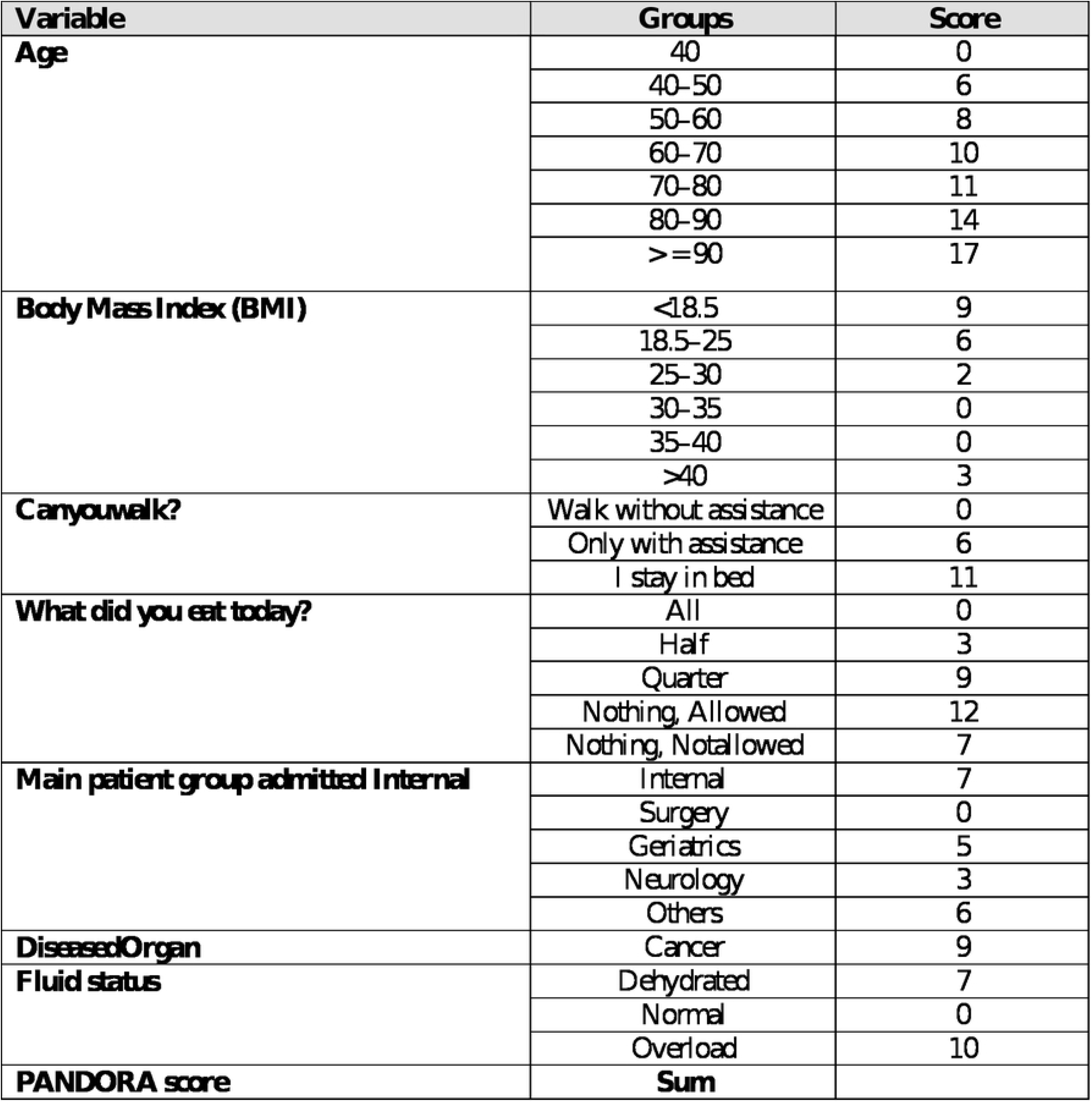
Original PANDORA Questionnaire

The applicability of the synthesis in Portuguese (2nd stage) was performed by means of paraphrase, in which the interviewer asks the question and asks the respondent to repeat it immediately. The synthesis in Portuguese was sent for retranslation, which was then compared with the original one to validate its Portuguese version.

### Data collection

Data collection took place from June to December 2018 and was carried out by previously trained nutritionists, using the instrument obtained from the final synthesis of PANDORA.

PANDORA is composed of 7 items related to the general evaluation with questions related to age, body mass index (BMI), physical activity level, hospitalizations, in-patient group, disease, hydration, and dietary assessment (amount of food ingested on the day of collection).

Each item quoted above generated a final score that was used to calculate the probability of death. The outcome (death or non-death) was followed up by searching the medical files of the hospital within 30 days of hospitalization.

### Statistical analysis

The collected data were stored in spreadsheets. A descriptive statistical analysis was performed, where the continuous data were presented in mean and average standard deviation. The normality of the data was tested by the Shapiro-Wilk test. In the presence of normality, an unpaired t-Student test was used to compare means. In the absence of normality, the U-Mann Whitney test was adopted. The relationship between the PANDORA score and hospital mortality at 30 days was assessed by the Logit formula (Logit = −6.72 + 0.1058 x PANDORA SCORE). From this result, the probability of death was calculated (e^logit^ / 1 + e^logit^).

Data on categorical variables were presented in absolute (n) and relative (%) values. Fisher’s exact test was performed to compare proportions between groups of categorical variables.

Logistic regression analysis (gross and adjusted) was performed using a death outcome to verify the association and the magnitude with the PANDORA score. From this analysis, the Odds ratio and its respective confidence interval were estimated. Finally, a ROC curve analysis was performed to evaluate the predictive power of the score on the outcome (death versus non-death) and estimated the area under the curve and its confidence interval.

The level of significance used for all tests was 5%. STATA® software version 12.0 was used.

## Results

The PANDORA questionnaire has been translated and adapted into Portuguese. No operational difficulties were observed during the paraphrase and retranslation test, allowing a reliable final Portuguese version (Fig. 3).

**Fig. 3.**
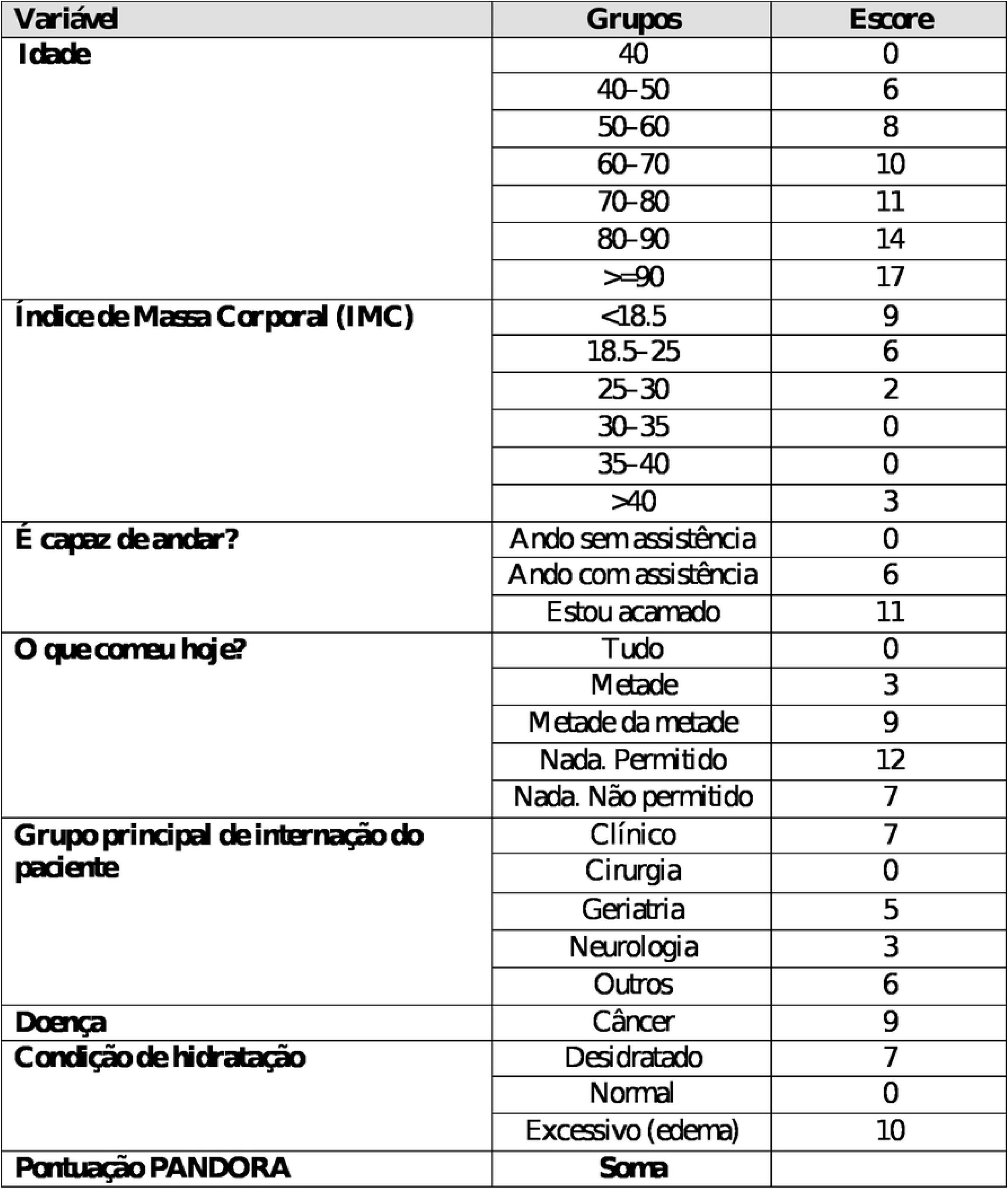
Portuguese version of the PANDORA questionnaire

The study included 54 patients with a mean age of 50.63 (sd = 16.81), of which more than half were male and 33% were elderly (> 60 years). More than 16% of the patients presented low weight, that is, BMI <18.5 kg / m^2^ for adults or <22 kg / m^2^ for the elderly, which was associated with death in the evaluated patients (p = 0.008). Among these low-weight patients, 12.5% reported that they did not eat anything on the day of data collection, 37.5% were fasting, 37.5% were 100% food acceptance, and 12.5% ate a quarter of what was offered on the day.

Considering the entire study population, the following data regarding food were observed: 28.8% of the patients ate half of what was offered; 6.6% did not eat anything; 12.9% were fasted; and 8.8% ate a quarter of the offer.

The mean PANDORA score was higher than 32 points in the general sample, however, with no significant difference in means in patients who died or not during hospitalization (p> 0.05). The prevalence of death in the study population was 9.2%. When evaluating the probability of death among patients, the mean was greater than 5% in the total sample (Table 1).

**Table 1.** Characterization of the study population and its relation with death / hospital discharge

| Variables | Total<br>n=54 | Death<br>n=5(9.26) | No death<br>n=49(90.74) | p-value |
| --- | --- | --- | --- | --- |
| **Age** | 50.63±16.81 | 48.6±9.29 | 50.84±17.44 | 0.780* |
| **Sex** |  |  |  |  |
| Male | 28(51.85) | 1(20.00) | 27(55.10) | 0.184** |
| Female | 26(48.15) | 4(80.00) | 22(44.90) |  |
| **BMI (kg/m<sup>2</sup>)** | 22.57±4.80 | 18.01±3.44 | 23.03±4.70 | **0.024\*** |
| Low weight | 9(16.67) | 4(80.00) | 5(10.20) | 0.008** |
| Eutrophy | 32(59.26) | 1(20.00) | 31(63.27) |  |
| Overweight | 10(20.41) | 0 | 10(20.41) |  |
| Obesity | 3(6.12) | 0 | 3(6.12) |  |
| **PANDORA score** | 32.13±9.66 | 37.20±11.75 | 31.61±9.41 | 0.221* |
| **Probability of death (%)** | 5.42±6.33 | 9.00±9.56 | 5.06±5.93 | 0.188‡ |
| **Classification of nutrition risk** |  |  |  | 0.144** |
| With risk | 33(61.11) | 5(100.00) | 28(57.14) |  |
| No risk | 21(38.86) | 0 | 21(42.86) |  |

Values presented in absolute values (relative values) or means ± standard deviation of the mean. p-value obtained by unpaired Student t-test or Fisher's exact test or Mann-Whitney test with 5% level of significance.

In the PANDORA questionnaire, only cancer disease is scored, with the other diseases classified as zero. It was verified that 83.3% (n = 45) of the studied population were diagnosed with some type of cancer, the most prevalent disease (79.62%), followed by Chronic Renal Disease, with a prevalence of 5.5%. Among the cancer patients, 17.7% had low weight, 100% presented nutritional risk. Of the ones who died, 100% were cancer patients.

In the adjusted logistic regression analysis, it was verified that there was no association between the PANDORA score and death in hospitalized patients; however, there was a trend of association of sex and BMI with death in these patients (Table 2). It was not possible to perform the analyzes with the probability of estimated death from PANDORA, due to the small sample size.

When verifying the predictive power of the PANDORA score on death in hospitalized patients using the ROC curve (Fig. 4), it was possible to verify an area under the curve of 0.66 considered adequate; however, when evaluating its range of confidence, we found that its lower limit was below 0.5, making the PANDORA score inadequate to predict death in patients in our sample.

**Fig 4.**
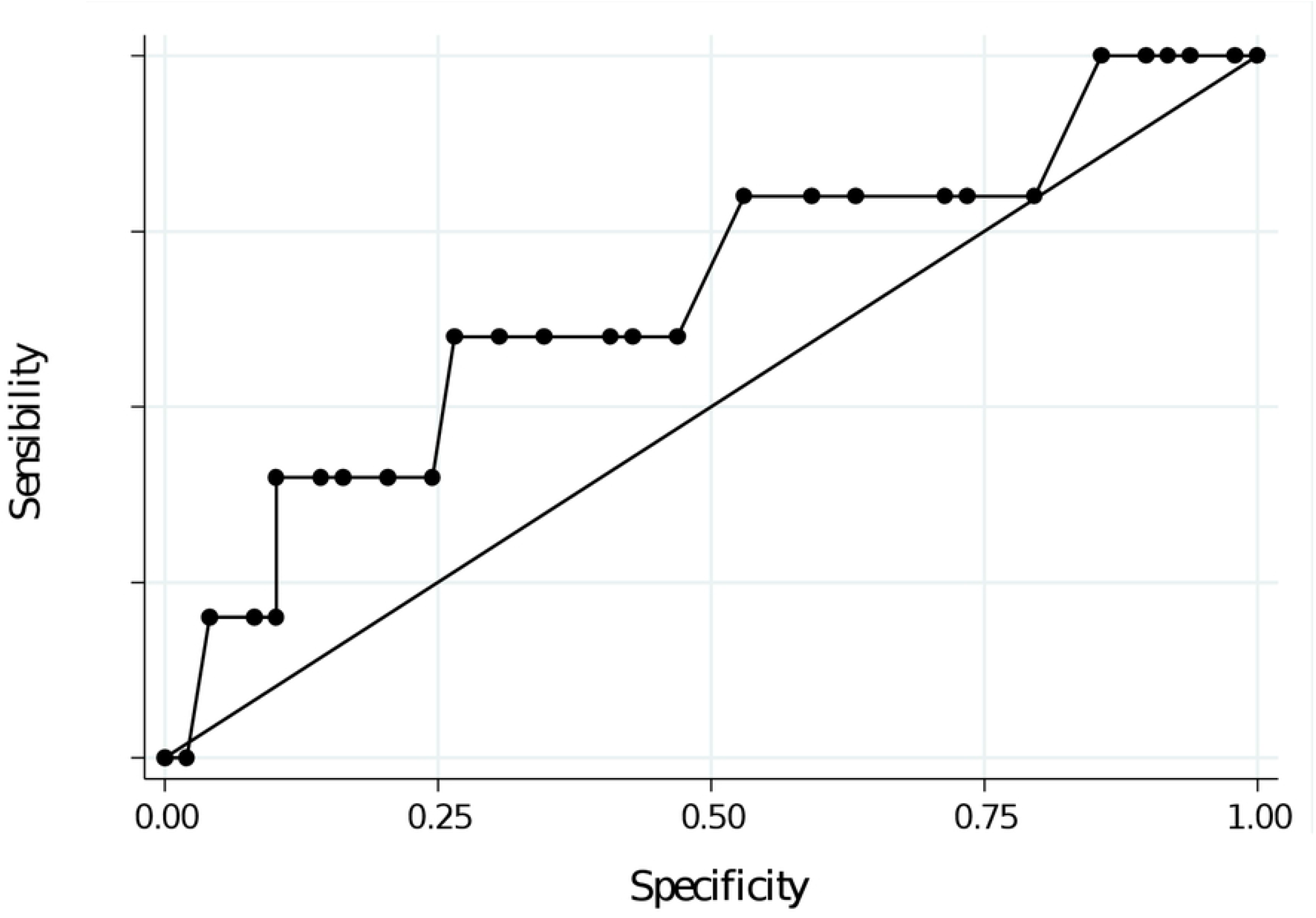
ROC curve

**Table 2.** Association among PANDORA score, demographic variables and nutritional status of the study population

|  | **Gross** |  | **Model 1** |  | **Model 2** |  | **Model 3** |  |
| --- | --- | --- | --- | --- | --- | --- | --- | --- |
|  | OR (IC95%) | p-value | OR (IC95%) | p-value | OR (IC95%) | p-value | OR (IC95%) | p-value |
| **PANDORA score** | 1.06(0.96-1.16) | 0.224 | 1.10(0.99-1.23) | 0.080 | 1.09(0.98-1.22) | 0.119 | 0.97(0.83-1.12) | 0.667 |
| **Age** | 0.99(0.94-1.05) | 0.775 | 0.95(0.88-1.03) | 0.219 | 0.95(0.87-1.03) | 0.206 | 0.98(0.90-1.08) | 0.754 |
| **Sex** |  |  |  |  |  |  |  |  |
| Male | 1.00 |  |  |  | 1.00 |  | 1.00 |  |
| Female | 4.91(0.51-47.16) | 0.168 |  |  | 4.78(0.45-51.17) | 0.196 | 28.97(0.98-851.81) | 0.051 |
| **BMI (kg/m<sup>2</sup>)** | 0.71(0.52-0.98) | 0.037 |  |  |  |  | 0.56(0.31-1.00) | 0.051 |

Logistic regression with Odds Ratio (OR) and 95% confidence interval (95% CI). Models: 1- Age-adjusted; 2 - Adjusted for age and sex; 3-Adjusted for age, sex, and BMI.

## Discussion

PANDORA has been recently developed and validated in a large multinational inpatient study, which included 2,480 patient care units in 32 countries. Its main advantage is the simplicity and practicality in the application to predict hospital mortality in 30 days.

In addition, it is based on nutritional markers, and may be useful for stratification of nutritional risk levels [11]. Another advantage found in the present study was the ease of translation and cross-cultural adaptation, since it is an instrument with direct questions and easy to understand.

The original PANDORA score had a high performance in predicting mortality in hospitalized patients having three main facilitator points: it is based on data available at any time of hospitalization; it is not necessary to spend much additional time collecting data, since the items are part of the patient’s history; and the model is public, international and independent of national codification conventions [2].

The association between low weight (BMI <18.5 kg / m^2^) and death in this study corroborates other studies found in the literature. Hu et al. (2017) evaluated the relationship between sarcopenic malnutrition syndrome and mortality in hospitalized elderly. It was observed that individuals with BMI between 18.6 kg / m2 and 19.5 kg / m2 were twice as likely to die compared with individuals whose BMI was greater than 22 kg / m2 [12]. Another study with elderly found similar results, and those with a mean BMI of 21 kg / m2 had a double risk of mortality [13].

A study conducted in the state of São Paulo, Brazil, observed that the presence of low weight was independently associated with higher mortality rates in patients who underwent coronary intervention [14]. Other authors performed a prospective nutritional screening and verified a strong association between mortality and risk of malnutrition, against our findings, in which 100% of the patients who died had nutritional risk [15].

In order to analyze the association between BMI and mortality in critically ill patients, a prospective multicenter study in France found that individuals with BMI <18.5 kg / m^2^ had an independent association with higher mortality. The authors also suggested that BMI may be a useful component in the development of future predictors of mortality. [16]. Tremblay and Bandi (2003) found similar results, with low weight being associated with a higher risk of mortality and worsening of functional status, delaying hospital discharge [17].

Hospital malnutrition should be evaluated very carefully since it is a public health problem, both in underdeveloped and in developing countries. The high prevalence of this problem persists over the years and presents worrying data [5]. In 1998 a study promoted by the Brazilian Society of Parenteral and Enteral Nutrition (BRASPEN) evaluated more than 4,000 hospitalized patients nationwide and found a prevalence of 48.1% of malnutrition [3]. In a recent systematic review, this high prevalence of malnutrition was confirmed [18]. All these findings reinforce the results found in our study, showing the importance of nutritional status in the outcome of hospital admissions [18].

Cancer is among the most common non-communicable diseases and injuries that cause death worldwide [19] and was the most prevalent disease in this study. According to the National Cancer Institute, the estimated incidence of the disease in 2018 was more than 300,000 new cases in Brazil [20]. Cancer patients are 1.7 times more likely to present malnutrition or nutritional risk than other hospitalized patients, both for physiological effects and for side effects to treatment [21]. A study of more than 2,000 cancer patients found that 19.7% of the individuals presented malnutrition, similar to our findings, where 17.7% of the cancer patients were underweight [22].

In this context, malnutrition in individuals with cancer is favored by low food intake, which was highlighted in this study, since most accepted only 50% of what was offered on the day. Ferreira, Guimarães and Marcandenti (2013) evaluated the acceptance of hospital diets of cancer patients admitted to a tertiary hospital and observed a high rate of rest ingestion among cancer patients and especially among the malnourished ones. The main associated factors were inappetence, xerostomia, constipation, dysgeusia, nausea related to smells and early satiety [23].

Another study based on Nutrition Day data collected between 2012 and 2015 evaluated the determinants of reduced food intake in colorectal cancer patients and concluded that being women, with advanced cancer, hospitalization longer than four days, and weight loss were the main determinants for the reduction of dietary intake of these patients. That study highlighted the need for the early identification of patients with nutritional risk for effective therapeutic measures [24].

Our study found no positive association between the PANDORA score and death, which makes it inadequate to predict death in the sample considered. This result can be justified by the limiting factor of the small sample size. The PANDORA score was already used in other studies and presented positive results as a tool to predict hospital death. Nakayama (2018), in an analytical cohort, evaluated whether the PANDORA score was associated with the mortality of patients in Intensive Care Units (ICU) compared with APACHE II. The authors concluded that the PANDORA score was strongly associated with mortality and that it can be compared to APACHE II for predicting mortality in critically ill patients [11].

Therefore, in the present study, the PANDORA score was not able to predict death in the patients in our sample, but found a significant association of low weight at admission with mortality.

## Conclusions

The PANDORA questionnaire was translated and adapted to Portuguese without any operational difficulties, allowing a final version that is reliable to the original one. It was not possible to predict death in patients in our sample using the PANDORA score. However, low weight had significant association with mortality, and may be an independent factor for predicting death.

In addition, a high prevalence of cancer in the studied population and association of the disease with the occurrence of low weight were observed, which highlights the need for studies that allow early identification of the nutritional risk factors in these patients, in order to obtain effective therapeutic plans.

Although some studies have shown positive results in relation to the PANDORA instrument, we suggest further studies for the Portuguese validation of PANDORA in hospitalized patients.

